# Uncovering the transcriptional molecular dynamics of shelf life extension and system acquired resistance induction to *Fusarium pallidoroseum* in melon fruits by the use of pulsed-light

**DOI:** 10.1101/2024.03.05.583617

**Authors:** Luis Willian Pacheco Arge, Guilherme Loss Morais, Joseane Biso Carvalho, Guilherme Julião Zocolo, Andréia Hansen Oster, Ana Tereza Ribeiro de Vasconcelos, Leandro Eugenio Cardamone Diniz, Ebenézer de Oliveira Silva, Patricia do Nascimento Bordallo

## Abstract

Melon is a globally commercialized fruit, and Fusarium rot disease poses a significant threat to post-harvest losses. The conventional use of fungicides raises concerns about chemical residues, prompting exploration into alternative technologies such as Pulsed Light (PL). While PL has been effective in controlling infections in various fruits and vegetables, the precise physiological responses and molecular mechanisms in melon fruits remain incompletely understood. In this study, melon fruits infected with the *Fusarium pallidoroseum* were treated with different doses of PL (0, 6, 9, and 12 J cm^−2^), and the impact on both fungal control and fruit shelf life extension was investigated. The 9 J cm^−2^ dose emerged as the most effective in controlling fungal growth without causing damage, inducing beneficial responses. This optimal PL dose upregulated genes in the lignan biosynthesis pathway and the infection upregulated genes involved with systemic acquired resistance, triggered by the pipecolic acid. In this way, the PL treatment and the infection trigger a double mechanism of resistance in melon fruits. A second and third experiment focused on evaluating the extension of melon fruit shelf life and the safe manipulation window post-PL treatment. The results revealed an average shelf life extension of six days and a safe manipulation period of 24 hours. The extension in shelf life was associated with a deviation in information flux from the ethylene biosynthesis to upregulation of the polyamine biosynthesis pathway, which produces nitric oxide, a product that can inhibit ethylene biosynthesis and its action. Furthermore, the observed 24-hour safety period against fungal infection post-PL treatment was characterized as a memory response resistance caused by the upregulation of lignan biosynthesis, which is a potential and efficient alternative to chemical products like fungicides. Overall, this study provides insights into the transcriptional molecular mechanisms through which PL promotes systemic acquired resistance and extends the shelf life of melon fruits.

## Introduction

In the years 2018 to 2022, the annual area planted with melon in Brazil averaged 25 thousand hectares (IBGE, 2023), corresponding to approximately 62 thousand acres or 250 square kilometers (km^2^) cultivated per year. This area produced an average of 622 thousand tons annually (IBGE, 2023). Of this total produced, Brazil exported around 233 thousand tons (37.5%) per year, generating export revenue of approximately 153 million dollars annually (COMEXSTAT, 2023). However, according to reports from Brazilian melon exporters (personal information), an estimated 15% of exported melons (approximately 35 thousand tons) exhibited symptoms of post-harvest diseases upon arrival at their destination and are consequently discarded. This results in a cumulative financial loss of approximately 23 million dollars. Yellow are the most melons exported by Brazil, but they are susceptible to pathogen attacks throughout the entire post-harvest chain. Among the main diseases affecting melons in the post-harvest stage, Fusarium rot stands out, caused by *Fusarium pallidoroseum* (Cooke) Sacc, synonym *Fusarium semitectum*. Despite the losses caused by fungal diseases, it’s worth noting that many fungi can produce metabolites that are toxic to human health (Oguro et al., 2015).

Today, consumers are more conscientious about the benefits of healthy and secure food, particularly with regard to chemical residues for pest and disease control. Countries importing melons have established rigorous limits for the maximum quantity of chemical residuals allowed. This has led to the rejection of containers due to the chemical concentrations exceeding the established limits or high losses caused by Fusarium rot disease. Consequently, there is a search for alternative technologies to control post-harvest diseases. In this context, one of the growing technologies to control post-harvest disease without chemical residuals is the use of physical treatments (Spadoni et al., 2015), such as pulsed light (Nascimento et al., 2014; Olaimat and Holley, 2012). This light, emitted in the range of 200 to 280 ηm, falls within the ultraviolet (UV-C) region and has germicidal effects (FDA, 2000; Bintsis et al., 2000) on a wide range of microorganisms (Rowan et al., 2015). These microorganisms include bacteria (Adhikari et al., 2015), protozoa, viruses (Hijnen et al., 2006), yeast (Niu et al., 2021) and fungi (Begum et al., 2009).

Ultraviolet light can be applied in a continuous or pulsed (UVP) manner. In UVP, the light is stored in a capacitor and released in intermittent flashes, resulting in a gain in radiant exposure over the time of application. This characteristic, combined with the number of applied pulses, makes it more effective in microorganism inactivation due to cellular membrane destruction (Elmnasser et al., 2007; Ignat et al., 2014). Additionally, UVP can stimulate the production of phytochemicals that protect plant tissues against the deleterious effects of radiation (Horev et al., 2012), and these phytochemicals can promote beneficial responses against other stresses, like biotic. This effect is known as hormesis, in which the application or treatment with small doses of harmful substances stimulate beneficial effects in an organism (Kendig et al., 2010; Agathokleous et al., 2019). In addition to the direct effect of UV-C, which is part of the UVP, on pathogen control, indirect benefits have been reported for fruits. These include the improvement of quality by eliciting specialized metabolic pathways (Darré et al., 2022) and extending fruit shelf life (Lu et al., 1991).

Plants employ various mechanisms to protect themselves against parasites and pathogens, leading to alteration in gene expression in both organisms (Benito et al., 1996). To better understand the mechanism of plant defense against phytopathogens, it is crucial to identify and characterize genes involved in the defense response. Advances in sequencing technologies, particularly automatic Sanger and Next Generation Sequencing (NGS), have revolutionized the discovery of new genes (Wang et al., 2009; Adams et al., 1991) and genes responsive to plant development (Schmid et al., 2005) as well as biotic and abiotic stresses (Coolen et al., 2016; do Amaral et al., 2016). Among these approaches, RNA-seq and its variations have emerged as the last most powerful techniques developed to date for studying the global gene expression of plants subjected to stresses. This method has been widely used to investigate post-harvest processes in various fruits, including strawberries (Hu et al., 2018), ioquat (Liu et al., 2019), blueberries (Zhang et al., 2020), watermelon (Guo et al., 2011), and more. In melon fruits, RNA-seq has been applied, for example, to study the association between carotenoid biosynthesis and mesocarp color (Diao et al., 2023), as well as the gene expression pattern of the melon-Fusarium pathosystem (Sebastiani et al., 2017).

A study of the transcriptional mechanism involving the dual interaction with the fruit, pathogen infection, and the effect of UV-C has not been reported yet. However, a recent study with *Musa nana* (M.-Z. Chen et al., 2021) described the transcriptional molecular mechanism triggering the defense response against diseases and the up-regulation of metabolite biosynthesis and antioxidant pathways through the use of UV-C. Our group’s previous study, involving metabolomics (Filho et al., 2020), serves as the foundation for this ongoing investigation to better understand the molecular mechanism behind the inducible resistance acquired and the extension of fruit shelf life promoted by UVP, using RNA-seq data. Our transcriptomics study has generated new insights into the molecular mechanism of the beneficial aspects promoted by the application of pulsed light in melons during the post-harvest stage.

## RESULTS

### Transcriptional profile analysis of melon fruits in response to pulsed light and the infection by *F. pallidoroseum*

To gain a better understanding of the transcriptional changes occurring in melon fruits infected with *F. pallidoroseum* and subjected to PL treatments (0 J cm^−2^, 6 J cm^−2^, 9 J cm^−2^ and 12 J cm^−2^), we conducted a comprehensive analysis of the melon fruit transcriptome using RNA-sequencing (RNA-seq) technology. This analysis provided new insights into the molecular mechanism underlying the response to Fusarium infection and to prolong the shelf life of melon fruits promoted by PL. Various comparisons of differential gene expression analysis were performed for this purpose.

Our control comparison included melons without infection and PL treatment (N00), melons infected and treated with 0 J cm^−2^ PL (I20), and melons without infection but treated with 9 J cm^−2^ of energy (N22). Additionally, we conducted other comparisons, such as the control without infection and treated with PL versus without PL (N22 vs N00), as well as comparisons within infections (I22 vs I21 and I23 vs I22). To access the quality control information for the libraries, please refer to supplemental table S1. The comparison of melon fruits infected with Fusarium (I20, I21, I22, and I23) against non-infected fruits (N00) revealed a total of 1788, 1302, 2098 and 1465 differentially expressed genes (DEGs), respectively. In the comparison of non-infected controls (N22 vs N00), a total of 1263 DEGs were identified. Non-infected fruits treated with PL of 9 J cm^−2^ (N22) were used as the base for comparing infected treatments against the same PL dosage or above (I22 and I23), resulting in a total of 809 and 414 DEGs, respectively. Furthermore, we compared the infected fruits, specifically those treated with PL (I21, I22, and I23), to those with 0 J cm^−2^ PL treatment (I20), revealing 62, 129 and 641 DEGs, respectively. Lastly, within the infection group, we found 7 and 462 DEGs for the I22 vs I21 and I23 vs I22 comparisons, respectively. All of these comparisons are depicted in figure S2, which includes their relationship through shared genes. Values corresponding to down- and up-regulated genes, as well as gene expression, are presented in supplemental table S2 and S3, respectively.

The number of DEGs alone does not explain the complex transcriptional systems biology behind the response to adverse conditions faced by the plants. Therefore, we performed some exploratory analysis using enrichment tests to uncover the main processes involved in the Fusarium attack on melon fruits and the effects of PL in controlling the infection and prolonging the shelf life. The analysis was carried out for all DEG profiles, separated into down- and up-regulated genes, using Gene Ontology (GO), KEGG pathways, and transcription factors annotations.

For GO, the enrichment analysis revealed a total of 368 overrepresented GO terms for biological processes across all profiles. Through the application of semantic similarity and dimensionality reduction, we were able to reduce this high complexity to just 19 main groups (Figure 1, Supplemental figure S3 and table S4). These groups are associated with light recognition/response encompassing red, blue and ultraviolet spectrum and light intensity; phenylpropanoid and flavonoid biosynthesis; lignin biosynthesis and catabolism; phenylalanine catabolism; processes involving abscisic acid responses; systemic acquired resistance mediated by the salicylic acid signaling pathway; response to ethylene; polyamine catabolic process; response to and metabolism of nitrogenous compounds; oxidation and reduction processes; and others.

**Figure 1.**
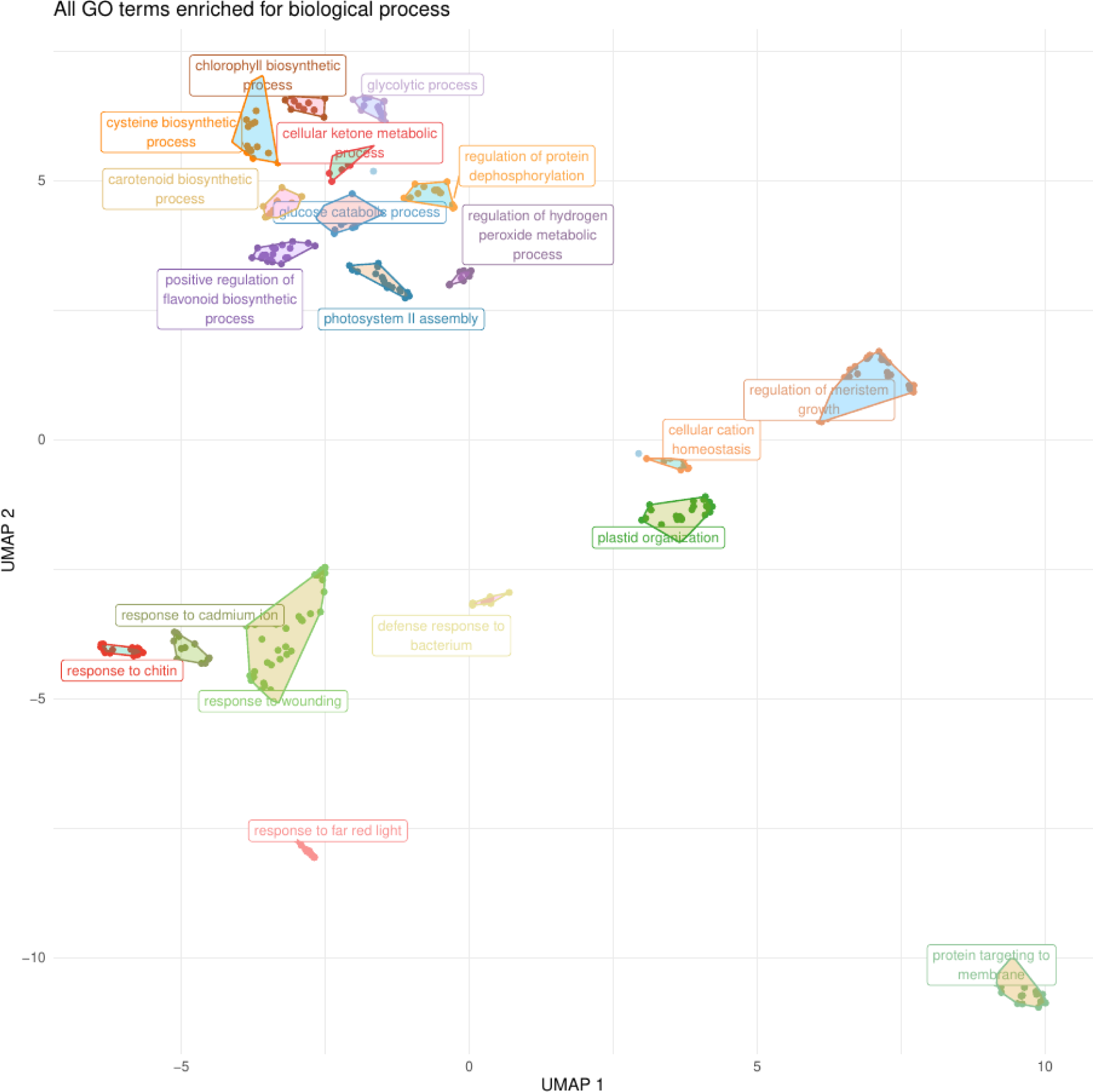
Overrepresented GO terms clustered for biological processes across all DEG profiles. Note: each cluster was labeled by the most significant GO term of the respective cluster.

The KEGG pathways enrichment analysis revealed that the biosynthesis of secondary metabolites, phenylalanine metabolism, phenylpropanoid biosynthesis, and photosynthesis (antenna proteins) were the main pathways overrepresented for the up-regulated genes. In total, only nine metabolic pathways were found across all DEG profiles out of a total of 25 (Figure 2A and Supplemental table S5).

**Figure 2.**
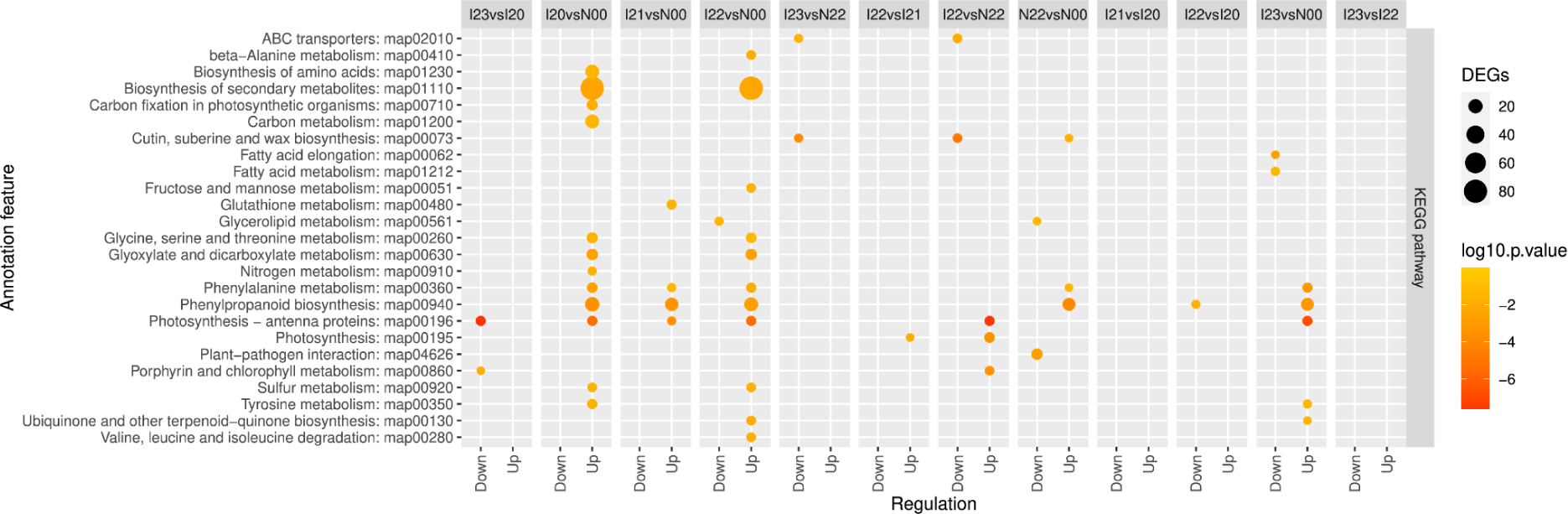
Enrichment analysis results for KEGG pathways from DEG profiles (up and down-regulated genes) of melon fruits infected by Fusarium and treated with pulsed light. The experimental design comprised six treatments, each with three biological replicates: N00: control (no inoculation, no PL treatment); I20: inoculation with Fusarium and treated with 0 J cm^−2^ PL; I21: inoculation and 6 J cm^−2^ PL; I22: inoculation and 9 J cm^−2^ PL; I23: inoculation and 12 J cm^−2^ PL; and, N22: without infection and treated with 9 J cm^−2^ PL. All PL applications were conducted 96 hours post-inoculation. The treatment nomenclature indicates whether there was no inoculation (N) or inoculation (I - first letter), the PL treatment time (0 or 2, indicating 0 hours or 96 hours post-inoculation - first number), and the PL dose applied (last number): 0, 1, 2, 3 (0, 6, 9, 12 J. cm^−2^, respectively).

Transcription factors (TFs) play a crucial role modulating gene expression. The enrichment analysis revealed a total of 2 enriched TF families, with WRKY (I21 vs N00 and I22 vs N22) and NF-YA (I21 vs N00) being the main families deregulated across all DEG profiles (Supplemental table S6).

### Information flux analysis of metabolic pathways through gene expression

Based on the results obtained from the enrichment analyses and a review of the literature, we hypothesized a flux of information across metabolic pathways (Figure 3) and this flux of information was investigated here, the enriched GO terms and metabolic pathways were cross-referenced and integrated through a literature review. The following results are described to provide profound insights into the effects of PL in controlling *F. pallidoroseum* infection and prolonging the shelf life of melon fruits.

**Figure 3.**
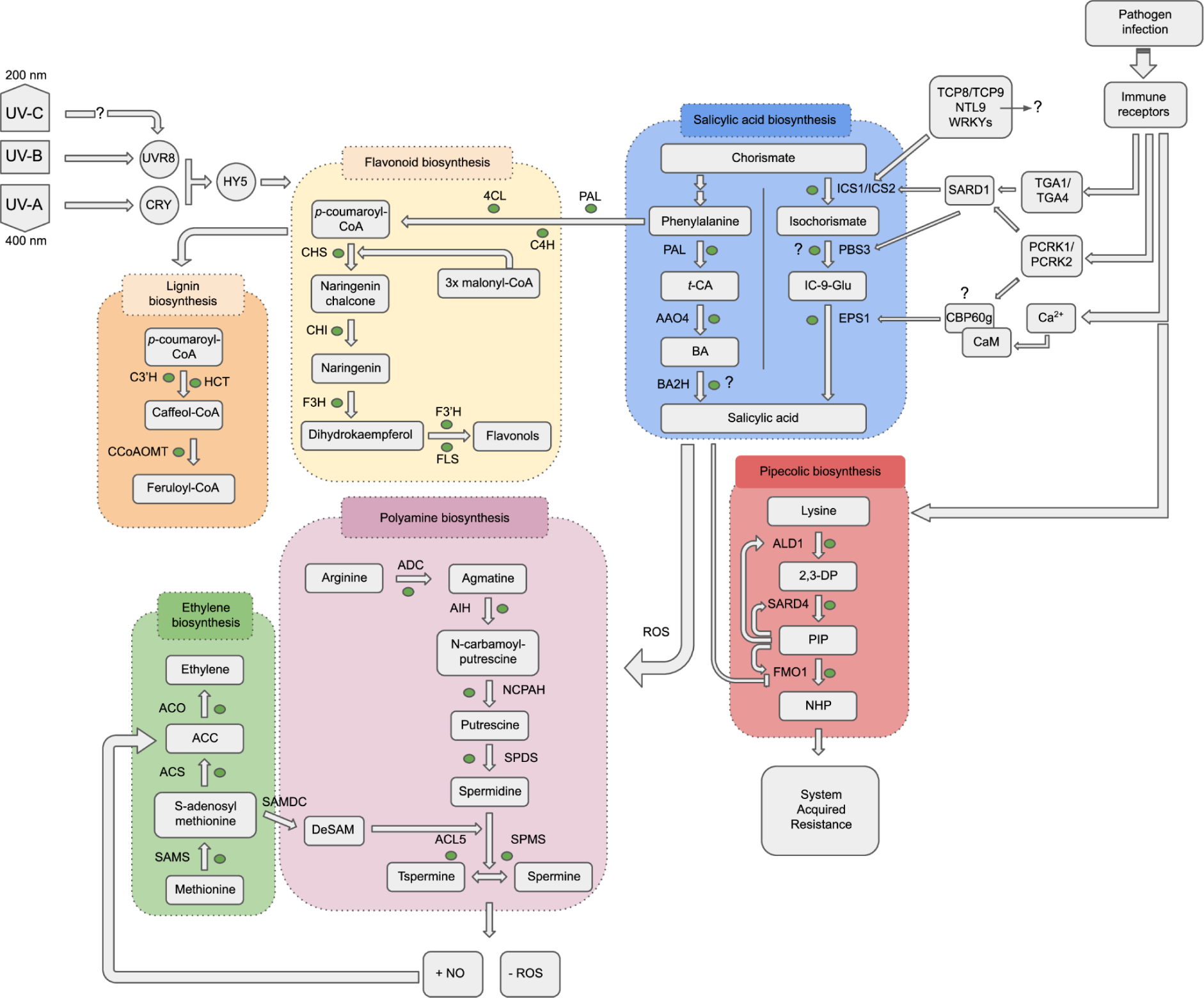
Reconstructed metabolic pathways to investigate the flux information triggered by pulsed light and Fusarium infection in melon fruits.

### Expression of photoreceptors genes in melon fruits treated with pulsed light

To evaluate our hypothesis model, we initially examined the transcriptional expression profile of known photoreceptors genes. These genes were identified through orthology analysis, comparing protein sequences from melon with arabidopsis and melon with tomato. Within the classical photoreceptors genes, we identified the expression of one *UVB-resistance 8* (*UVR8*) gene and three *cryptochromes* (*CRY*) genes. Among them, only one *CRY*, specially the copy three (MELO3C003644), showed differential expression, being down-regulated in the N22 vs N00 comparison (Figure 4A). However, other genes participate in the cascade of light recognition signals, such as *hypersenescence 1* (*HYS*) and *constitutive photomorphogenic 1* (*COP1*), two transcription factors that showed expression but not differential expression for the comparisons.

**Figure 4.**
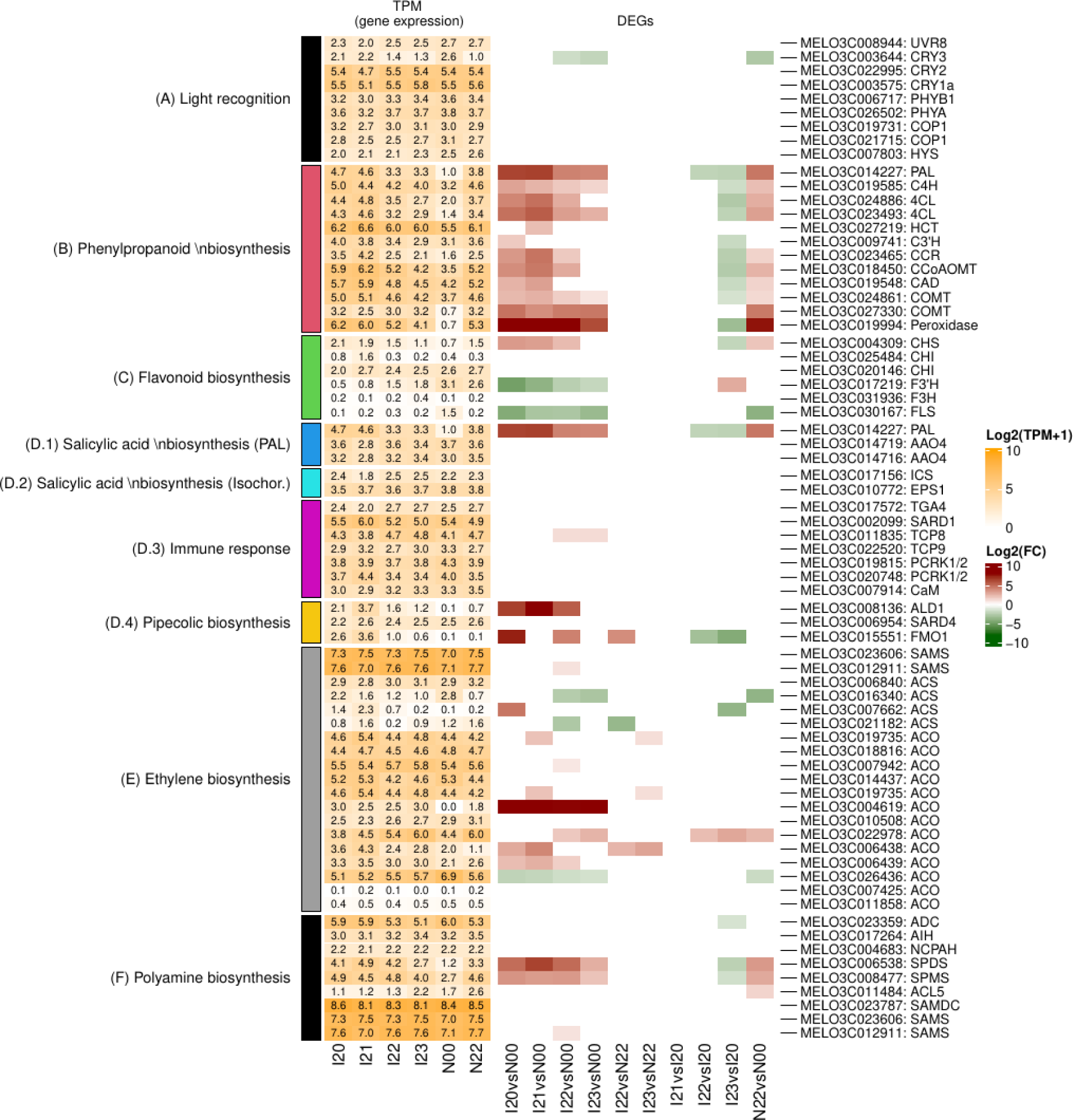
Gene expression and differential expression values for genes involved with biological pathways responsive to pulsed light in melon fruits. The experimental design comprised six treatments, each with three biological replicates: N00: control (no inoculation, no PL treatment); I20: inoculation with Fusarium and treated with 0 J cm^−2^ PL; I21: inoculation and 6 J cm^−2^ PL; I22: inoculation and 9 J cm^−2^ PL; I23: inoculation and 12 J cm^−2^ PL; and, N22: without infection and treated with 9 J cm^−2^ PL. All PL applications were conducted 96 hours post-inoculation. The treatment nomenclature indicates whether there was no inoculation (N) or inoculation (I - first letter), the PL treatment time (0 or 2, indicating 0 hours or 96 hours post-inoculation - first number), and the PL dose applied (last number): 0, 1, 2, 3 (0, 6, 9, 12 J. cm^−2^, respectively).

Furthermore, we expanded our analysis to a set of 710 genes [which included at least one gene with TPM (Transcript Per Million) ≥ 1 across the DEG profiles] annotated with the enriched GO terms related to light responses. Within this gene set, we identified several genes that are indirectly involved in light recognition. Specifically, we detected 11 genes related to peroxidase, 12 genes related to phenylalanine ammonia-lyase (PAL), 24 genes related to phytohormones (such as auxin, abscisic acid, ethylene, and gibberellin), and 41 genes related to zinc finger and/or transcription factors. Moreover, we discovered 190 genes that have not been functionally attributed based on homology detection (Supplemental table S3). This finding, of uncharacterized genes, presents an opportunity to investigate potential new genes involved in light responses and possibly UV-C recognition.

### Phenylpropanoid and flavonoid biosynthetic gene expression

One of the frequently associated pathways triggered by pathogen attack and light sensing is the phenylpropanoid and flavonoid biosynthetic pathways. Using orthology identification from Arabidopsis and tomato, we were able to reconstruct these pathways in melon species (*Cucumis melo*). We detected 12 genes with differential expression in the phenylpropanoid pathway, showing a clear up-regulation pattern in the comparisons of infected conditions (I2X) against control (N00). However, a down-regulation pattern was observed in the comparison of I23 vs I20, and no DEGs were found in the I22 vs N22, I23 vs N22, and I21 vs I20 profiles (Figure 4B).

In the flavonoid biosynthesis pathway, we identified six ortholog genes, including the gene *chalcone synthase* (*CHS* - MELO3C004309) from the initial pathway. This gene was detected as up-regulated in the I20, I21 and I22 vs N00, as well as N22 vs N00 DEG profiles. However, the majority of the genes in the pathway showed down-regulation or only exhibited gene expression patterns without significant changes (Figure 4C). This indicates an information flux pattern towards the phenylpropanoid biosynthesis pathway (Figure 4BC), suggesting an up-regulation of lignins instead of flavonoid biosynthesis.

### Salicylic and pipecolic acids biosynthesis as mediator of system acquired resistance

Salicylic acid is synthesized from the chorismate molecule and, currently, there are two possible pathways to salicylic acid biosynthesis in plants. One pathway is derived from phenylalanine, which is also a precursor for phenylpropanoid and flavonoid molecules, and the other pathway is derived from isochorismate. In our analysis, we identified only one DEG for each of these chorismate-derived pathways, the *phenylalanine ammonia lyase* (*PAL*) (Figure 4D.1). Additionally, we detected the expression of two *aldehyde oxidase 4* (*AAO4*) copies in the phenylalanine-derived pathway and two genes, *isochorismate synthase* and *enhanced pseudomonas susceptibility 1* (*ICS* and *EPS1*, respectively), in the isochorismate-derived pathway (Figure 4D.2).

Genes associated with the immune response are known to be involved in the biosynthesis of salicylic acid. In our study, we observed differential expression for three out of seven immune response genes. These genes include *SAR deficient 1* (*SARD1*), *TCP domain 8* (*TCP8*) and *pattern-triggered immunity compromised receptor-like cytoplasmic kinase 1/2* (*PCRK1/2*). However, all of the non-DEG gene copies were detected as expressed across all gene expression profiles (Figure 4D.3).

Salicylic acid triggers the system acquired resistance by acting as a modulator for pipecolic acid biosynthesis. In our analysis, we identified all the three orthologous genes involved in this pathway. *L-lysine alpha-aminotransferase 1* (*ALD1*) and *flavin-dependent monooxygenase 1* (*FMO1*) genes were found to be differentially expressed, while *SAR deficient 4* (*SARD4*) showed similar expression values across all conditions and no differential expression (Figure 4D.4).

### Expression of ethylene and polyamine biosynthetic genes

Ethylene is one of the primary phytohormones involved in fruit maturation, and this pathway can be diverted towards the biosynthesis of polyamines. In our analysis, we identified a bias in the information flux towards the biosynthesis of polyamines despite ethylene (Figure 4E and F). We identified two gene copies for *S-adenosyl-l-methionine* (*SAMS*), with TPM expression ranging from 7.0 to 7.6 and one gene (MELO3C012911) showed differential expression in the I22 vs N00 profile (Figure 4E). The next genes in the pathways are *1-Aminocyclopropane-1-carboxylic acid synthase* (*ACS*), which is involved in ethylene biosynthesis, and *S-adenosylmethionine decarboxylase* (*SAMDC*), which is involved in polyamine biosynthesis (Figure 4F). We observed that the expression of *SAMDC* is significantly higher than that of all four copies of *ACS*. For *1-aminocyclopropane-1-carboxylic acid oxidase* genes (*ACO*), we identified 13 gene copies expressed across all profiles, and some of these showed differential expression. Lastly, the polyamine biosynthetic pathway is composed of seven genes, for which orthologous copies were identified and their expression was detected. Interestingly, the genes *spermidine synthase* (*SPDS*), *spermine synthase* (*SPMS*) and *thermospermine synthase* (also known as *acaulis 5* - *ACL5*) from the end of the pathway were found to be upregulated in the N22 vs N00 comparison, as well as in the I20, I21, I22 and I23 vs N00 comparisons (Figure 4F).

### Phenotypic evaluation of infection post pulsed light application

Goldex^Ⓡ^ melon fruits were evaluated in terms of the treatment of pulsed light (PL) to control the infection caused by the *F. pallidoroseum* and to extend the fruit shelf life. Initially, the melon fruits were submitted to the application of different doses of PL: 0, 9 and 12 J cm^−2^ and the infection with Fusarium was performed at 0, 24, 48 and 96 hours after the light treatment (Figure 5A), and then the fruits were stored at conditions of 5-7°C and 80% (±5%) of relative humidity. The growth of the infection diameter was monitored over 18 days in the different treatments (Figure 5B). Comparing the treatments we discovered the application of PL light of 9 J cm^−2^ was the best treatment to control the Fusarium infection occurring immediately or until 24 hours after PL application (Figure 5AC). 12 J cm^−2^ treatment presented the highest damage to the melon fruit tissues as observed in the first experiment, showing cell-wall degradation and loss of firmness. However, a recovery period was observed from 0h to 24h in 12 J cm^−2^, and a similar pattern was observed in the infection occurring after 96h of fungicide application independent of pulsed light treatment. Additionally, a residual physical effect of PL was observed and this residual acts controlling the Fusarium infection in melon fruits.

**Figure 5.**
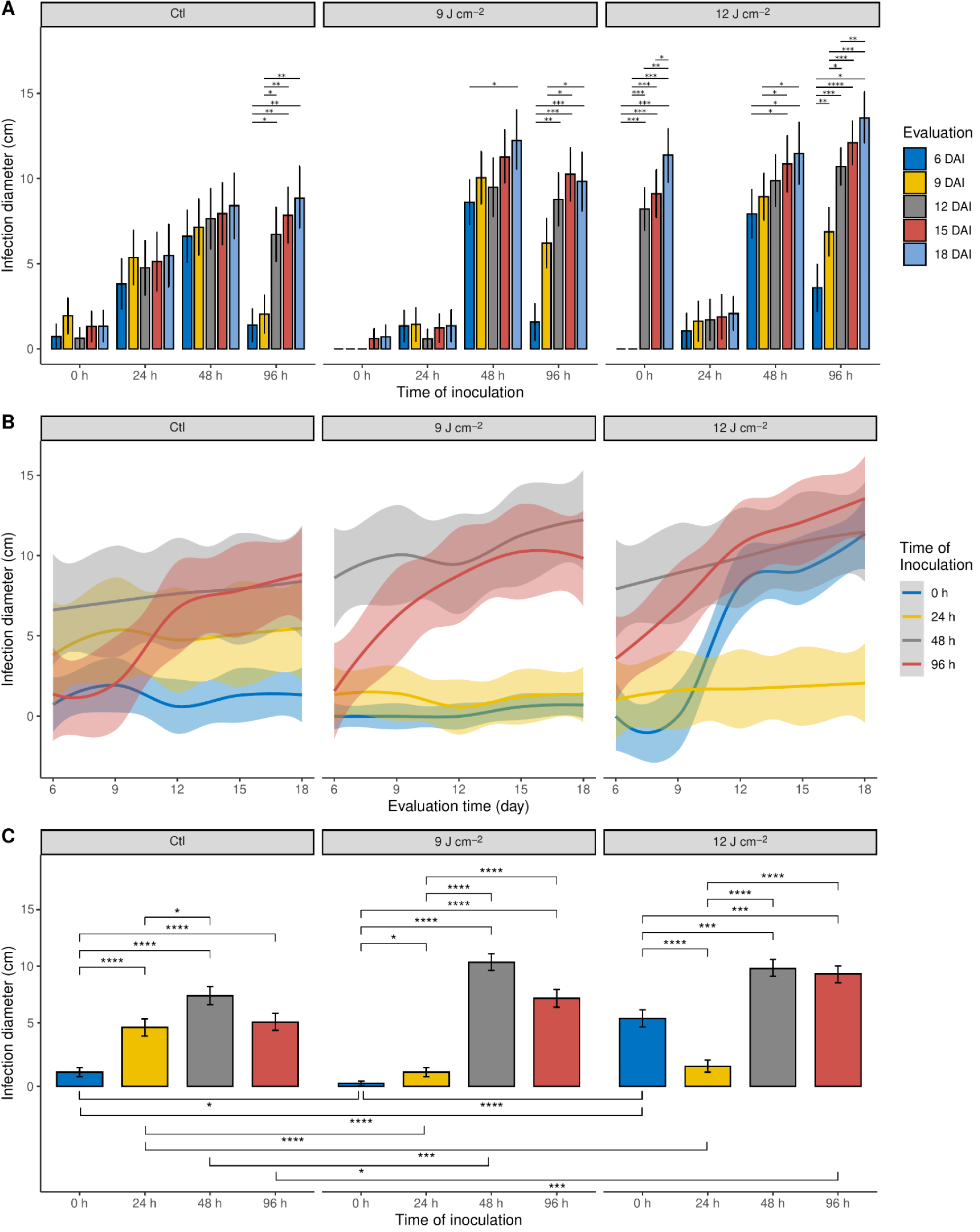
Phenotypic evaluation of melon fruits treated with pulsed light doses (0, 9 and 12 J cm^−2^) to control *Fusarium pallidoroseum*. Subfigure A shows the infection diameter on melon fruits in different times and days, and subfigure B shows the loess regression (a non-parametric approach) with a fitted smooth curve for the different times of inoculation across the days. Note: * < 0.05, ** < 0.01, *** < 0.001 and **** < 0.0001, and vertical lines over the bars represent standard error.

Regarding the shelf life of the melon fruits, a survival analysis was conducted, which revealed a median survival probability of 23 days for fruits without PL treatment and 29 days for melon fruits treated with PL (Figure 6), resulting in a gain of 6 days. Within 23 days, the median observed survival count for untreated fruits was 69. At this point, it was noticeable that 33 fruits had decayed. Conversely, for PL-treated fruits, the median survival count was 58 at 29 days, meaning that more than half of the fruits reached 29 days with PL treatment.

**Figure 6.**
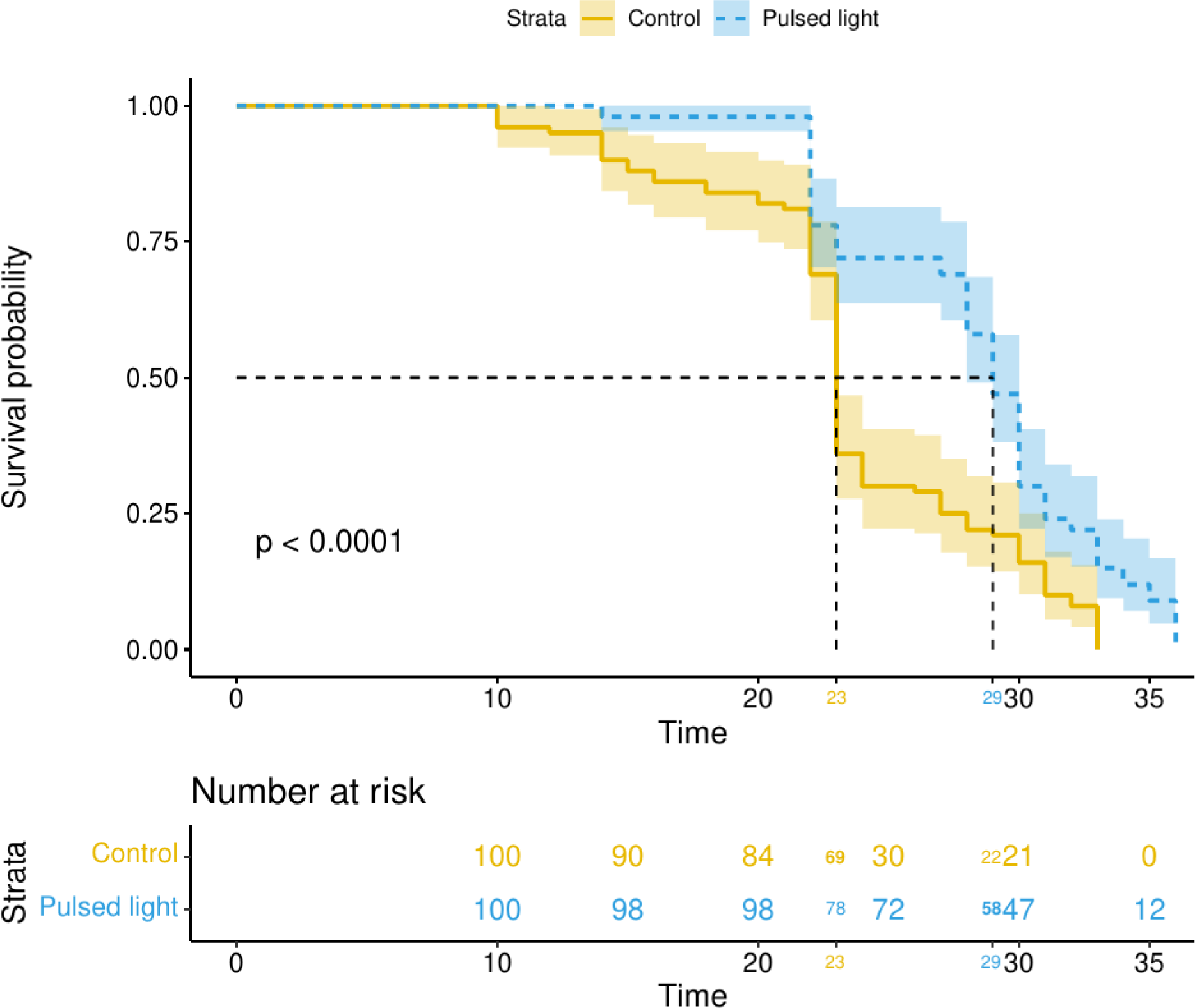
Survival distribution curves of melon fruits treated with pulsed light (9 J cm^−2^), and controls (0J), both conditions without Fusarium inoculation. Evaluation of shelf life was performed during 35 days under conditions of 5 - 7°C and 80% (+/−5%) of relative humidity (RH).

## Discussion

### Transcriptional Dynamics of Melon Fruits Subjected to Pulsed Light and Infection by the *F. pallidoroseum*

Pulsed light technology has emerged as a promising method for controlling fungal infections in various fruits during the post-harvest stage. In this context, our study represents a logical continuation of our prior research, which utilized metabolomics technology as detailed in (Filho et al., 2020). The previous study employed the same experimental design to examine the metabolic changes occurring in melon fruits. However, the current research sought to delve deeper and unravel the transcriptional mechanisms responsible for the effects of PL on controlling the Fusarium infection and the pathways associated with post-harvest preservation in melon fruits. The results obtained from the transcriptional analysis serve to not only validate our prior findings in metabolomics and phenotypic data but also to provide a more comprehensive understanding of the intricate molecular mechanisms that underpin the effects of PL to enhance quality and fungal resistance, especially in the post-harvest management of fruits.

Our differential gene expression analysis has unveiled that the response to PL is a holistic process, involving the deregulation of various biosynthesis pathways. These pathways encompass those associated with phenylpropanoids, salicylic and pipecolic acids, ethylene, and polyamines. Furthermore, this deregulation has downstream effects on oxidative-reduction processes and the induction of systemic acquired resistance. In this context, previous studies conducted by our research group have produced findings that align with the results of the present study. For instance, Sousa et al. (2019) observed similar deregulation in the ethylene and polyamines metabolic pathways after treating melon fruits with PL. However, other studies have also explored the transcriptional molecular mechanisms underlying the sanitation effects of PL. One such study, conducted by Wang et al. (2023), investigated the impact of PL on controlling *Aspergillus carbonarius* in pear fruits. Their research revealed that a high dose of PL inhibited mycelial growth by 30% with a PL dose of 13.5 J cm^−2^ and significantly reduced ergosterol and ochratoxin A (OTA) content by 27.9% and 80.7%, respectively. Ergosterol is a crucial component of fungal cell membranes and is a target for the development of antimicrobial agents (Kumar and Jha, 2023). OTA is an important toxin associated with food safety concerns (Alfonso et al., 2022), so these results have significant implications for fruit quality and safety. Furthermore, the transcriptome analysis of the fungus revealed the down-regulation of genes associated with cell integrity, energy and glucose metabolisms. These findings align with their results of cellular leakage observed through transmission electron microscopy, and these results can be complementary to our results on revealing the molecular fungal mechanism of control promoted by the PL.

It’s indeed noteworthy that previous studies outside our research group have not extensively explored the molecular effects of PL on controlling fungal infections and extending fruit shelf life. Our study bridges this gap by providing insights into the transcriptional mechanism involved in these processes. The transcriptional expression results of our study align with the metabolomics findings from (Filho et al., 2020), particularly regarding the specialized metabolic pathway of phenylpropanoids. This pathway emerges as a crucial complex adaptive mechanism in response to both biotic and abiotic stressors. These findings also find support in the results from Lopes et al. (2017), who observed alterations in the phenylalanine metabolism and the biosynthesis of phenylpropanoids as significant outcomes in mangoes treated with PL. These metabolic changes, coupled with the induction of acquired resistance, seems to represent the primary cellular processes in the fight against fungal infections.

Transcription factors play a central role in modulating the exchanges within these processes. The enriched families of transcription factors identified are diverse and play important roles in both abiotic and biotic stresses. WRKY is recognized as an army of stress-fighting soldiers (Viana et al., 2018). However, while many transcription factor families present vital roles in abiotic and biotic stress, few of these have been intensively studied, as the case of NY-FA. It is important to mention that one or few transcription factors of a family can play important roles in specific biological processes, resulting in phenotype changes. The findings of Sebastiani et al. (2017), who studied a different melon cultivar infected with *Fusarium oxysporum*, corroborate with our findings on WRKY transcription factor family. Thus, they reinforce the importance of the WRKY family in orchestrating responses to fungal infections.

The exploration of the effect of PL on sanitation is well-established, but its impact on the shelf life of fruits is less understood. Through the development of previous studies with PL on post-harvest fruit treatment in our group, we noticed an extension in the shelf life of fruits. This extension was also recently reported in banana fruits treated with UV-C (Chen et al., 2021), where an accumulation of specialized metabolites associated with fruit quality was observed, and since at least 1991 for many fruits (Lu et al., 1991). Similarly, high intensity light has been reported to increase fruit shelf life in many vegetables (Poiroux-Gonord et al., 2010).

In our study, utilizing transcriptomics and metabolomics data, we delved into the molecular mechanisms underlying the extension of fruit shelf life alongside pathogen control. In the context of fruit shelf life extension, genetic reprogramming emerges as a more crucial factor than metabolic processes, when compared with pathogen control or resistance. Leveraging a highly annotated genome with Gene Ontology (GO) terms and employing clustering to identify groups of highly similar enriched GO terms, along with the analysis of metabolic pathways, enabled the reconstruction of pathways associated with fruit conservation and systemic acquired resistance. The analysis of information flux through these pathways offered a better comprehension of a complex molecular mechanism, and this is discussed in detail below.

### Perception of pulsed light effects seems to not be by the classical photoreceptors, but a holistic perception

Light stimuli in plants are perceived by specific receptors known as photoreceptors. Among these, cryptochromes recognize UV-A, and UV-B is recognized by UV-B resistance 8 proteins (Paik and Huq, 2019). This recognition initiates a cascade of signal transduction events leading to molecular changes. However, receptors for UV-C have not been fully characterized yet (Rai et al., 2021), presenting a bottleneck in understanding the flux to molecular changes. In our study, almost all orthologs of photoreceptors did not show differential expression, except for CRY3, which was down-regulated.

From these results, we hypothesize that pulsed light might be recognized by various cellular molecules due to its range of energy content, eliciting different molecules, proteins and enzymes that could lead to metabolic changes. An intriguing aspect is that PL cannot penetrate materials deeply (Meinhardt et al., 2008), and due to the epidermis differences of the fruits the PL dose should be adjusted to each one in order to promote beneficial responses. Melon fruits, with a thicker superficial barrier compared to strawberries, may require more energy to trigger beneficial responses related to induced resistance and post-harvest shelf life extension, instead of damages.

### A physical barrier stimulated by PL through the lignan biosynthesis pathway

The lignan and flavonoid biosynthesis pathways are recognized for producing compounds with antimicrobial and antioxidant properties, which can act as UV protectants (Tohge et al., 2017). The regulation of flavonoid biosynthesis is well known to be modulated by UVR8 (Clayton et al., 2018), which, in our study, did not show differential expression. Consequently, most genes in the flavonoid biosynthesis pathway exhibited no differential expression or were down-regulated. An exception was *chalcone synthase* (*CHS*), which encodes an enzyme that converts *p*-curamoyl-CoA to naringenin chalcone, a metabolite detected in high abundance in the (Filho et al., 2020) study, and also *p*-curamoyl-CoA is a precursor for salicylic acid biosynthesis.

Both flavonoid and lignan biosynthesis pathways derive from phenylpropanoids. The information flux analysis results suggest a deviation from flavonoid to lignan biosynthesis, indicated by the down-regulation of flavonoid genes and the upregulation of lignan genes. Lignan, as the second most abundant biopolymer in plants, constituting 30% of organic carbon, is known to enhance plant defense during biotic stresses (Lee et al., 2019). It improves cell wall rigidity, establishing a physical barrier against fungal and other infections. This aligns with our indirect phenotypic observation of increased firmness in fruits treated with a 9 J cm^−2^ PL dose and their enhanced resistance to fungal infection.

### Pipecolic acid as a trigger for systemic acquired resistance induction

Despite the deviation of information flux towards the biosynthesis of lignans, known genes in the salicylic acid biosynthesis pathway showed expression, although no significant differential expression was observed. While the full set of genes for the two potential pathways of salicylic acid biosynthesis in melon has not been fully characterized, the expression of orthologous genes was detected for some of these genes, indicating activity of this pathway. Similarly, orthologous genes related to immune response were identified, displaying global expression without significant differential expression.

Intriguingly, downstream genes in the salicylic acid pathway, specially those involved in the biosynthesis of pipecolic acid (PIP), exhibited differential expression. Pipecolic acid is recognized as a trigger for systemic acquired resistance in plants (Bernsdorff et al., 2016). The mechanism involves salicylic blocking the activity of Flavin-Dependent-Monooxygenase1 (FMO1), an enzyme that converts PIP into N-hydroxy-pipecolic acid (NHP). However, when PIP concentration is high, it promotes FMO1 activity bypassing the blocking effect promoted by salicylic acid, catalyzing in this way the conversion of PIP into NHP. Additionally, the high concentration of PIP stimulates the conversion of lysine into Δ^2^-piperideine-2-carboxylic acid (2,3-DP) by promoting the activity of the AGD2-Like Defense Response Protein 1 (ALD1) enzyme, which is the immediate precursor metabolite of PIP, and also stimulates the conversion of 2,3-DP into PIP by the Systemic Acquired Resistance-Deficient 4 (SARD4) enzyme. This is an elegant regulation mechanism of a biosynthesis pathway, whose product is the NHP, a molecule that triggers a local and a systemic acquired resistance via transport to adjacent parts in the plant (Chen et al., 2018; Bernsdorff et al., 2016).

### Extension of shelf fruit life seems to be by a flux deviation from the ethylene biosynthesis to the polyamine biosynthetic pathway

The primary pathway investigated during the post-harvest stage to prolongs the fruit shelf life is the biosynthesis of ethylene, a gas hormone crucial for the maturation process (Iqbal et al., 2017). Chemical products like 2-aminoethoxyvinyl glycine (AVG), silver ions (Ag), and the gaseous compound 1-methylcyclopropene (1-MCP) have been used to inhibit ethylene action in fruits, successfully extending shelf life (Schaller and Binder, 2017). However, these chemical products may leave residuals affecting health, and genetic reprogramming to control ethylene biosynthesis requires government permission for a restricted commercialization. UV-C application has been linked to extending the shelf life of fruits since at least 1991 (Lu et al., 1991). Pulsed light, which includes UV-C wavelengths, promotes shelf life extension in melon fruits, and this process may be explained by the content of UV-C presented in PL. However, the mechanism behind this beneficial effect is not fully understood.

With the information flux analysis results, a deviation from the early ethylene biosynthesis pathway to the biosynthesis of polyamines was observed. The enzyme S-adenosyl-l-methionine synthase is encoded by the gene *SAMS*, which showed expression across the conditions. Interestingly, in the polyamine biosynthesis pathway, *S-adenosyl-l-methionine decarboxylase* (*SAMDC*), a gene deviating towards polyamine biosynthesis, exhibited higher expression than the following ethylene-related gene, *1-aminocyclopropane-1 carboxylic acid synthase* (ACS). In a previous study of our group, was constated inhibition of ethylene biosynthesis and ACC synthase and ACC oxidase activities remained unaltered on melon during storage at 4°C [PL alone and PL +1-MCP, Sousa et al. (2019)].

Further along the polyamine biosynthesis pathway, all genes presented expression. Notably, the end genes, *spermidine synthase* (*SPDS*) and *spermine synthase* (*SPMS*), showed differential expression values in tissues infected with Fusarium, with or without PL treatment. Control comparisons with PL without infection also displayed differential expression for these genes, along with another polyamine gene, *acaulis 5* [*ACL5*, Tun et al., (2006)]. These genes catalyze the conversion of putrescine to spermidine (*SPDS*) and then to spermine (*SPMS*), and spermidine to thermospermine (*ACL5*), which means a basal expression of this pathway in melon fruits treated with PL.

Spermidine has been used to delay the ripening of peaches, and its effect seems to be mediated by the polyamine metabolism subproduct nitric oxide [NO, Gupta et al., (2022)]. NO acts as transcriptional repressor of genes involved in ethylene biosynthesis, reduces methyl groups required for ethylene production, and competes with ethylene receptors (Manjunatha et al., 2012; Gupta et al., 2022), altering the expression of genes related to cell wall metabolism and lignification, as observed in this study. Based on this, the biosynthesis of polyamine seems to be the main pathway responsible for extending the melon fruit shelf life.

### Pulsed light induces Fusarium resistance memory and prolongs the shelf life of melon fruits, the phenotypic evaluation

The first experiment of this study was used to uncover the transcriptional molecular mechanism of PL effect on controlling the fungal growth on melon fruits already infected. This effect of superficial and low depth sanitation had been reported by several studies on fruits, including strawberries, cherries, apples, raspberries, grapes, nectarines, peaches, pears, lemons, oranges, kiwis, mangoes, and melons (Lagunas-Solar et al., 2006; Sebastiani et al., 2017; Filho et al., 2020; Lopes et al., 2017; Sousa et al., 2019; Marquenie et al., 2002; Contigiani et al., 2021). However, no one reported the post effect of PL on controlling fungal growth, a hypothesis that was raised on our group through several previous experiments, which could be an alternative technology to fungicide applications to control fungal infections, mainly the infections caused by Fusarium that promotes several losses in this culture.

Based on the transcript-/metabol-omics experiment we could determine the 9 J cm^−2^ PL dose as the most effective to control fungal infection in melon fruits (Supplemental figure S4), dose which was used together with control and 12 J cm^−2^ to evaluate the residual, or memory, effect of PL on controlling post harvesting infections. By the phenotyping experiment we could validate our hypothesis of memory resistance triggered by PL, specifically for the 9 J cm^−2^ dose, the same dose mentioned above for local sanitation. This dose was particularly most effective for controlling infections in a period of 24 h after the PL application. Especially, a period of 12 h after the harvesting is crucial to control post-harvesting disease, because in these first hours occurs the manipulation of the fruits, when they can be infected by post-harvest fungal microorganisms responsible for diseases which promotes up to 30% of losses in melon fruit culture. Our results revealed a safety period of manipulation promoted by PL that is more than enough to avoid important post-harvest infections in the melon fruit culture, and this technology can be used as an alternative to the agrochemical agents, consequently promoting a better food security (Figure 7). However, future studies will be needed to reveal if this memory resistance is triggered by the transcriptional modulation resulting in the up-regulation of lignans genes or by the accumulation of other specific metabolites.

**Figure 7.**
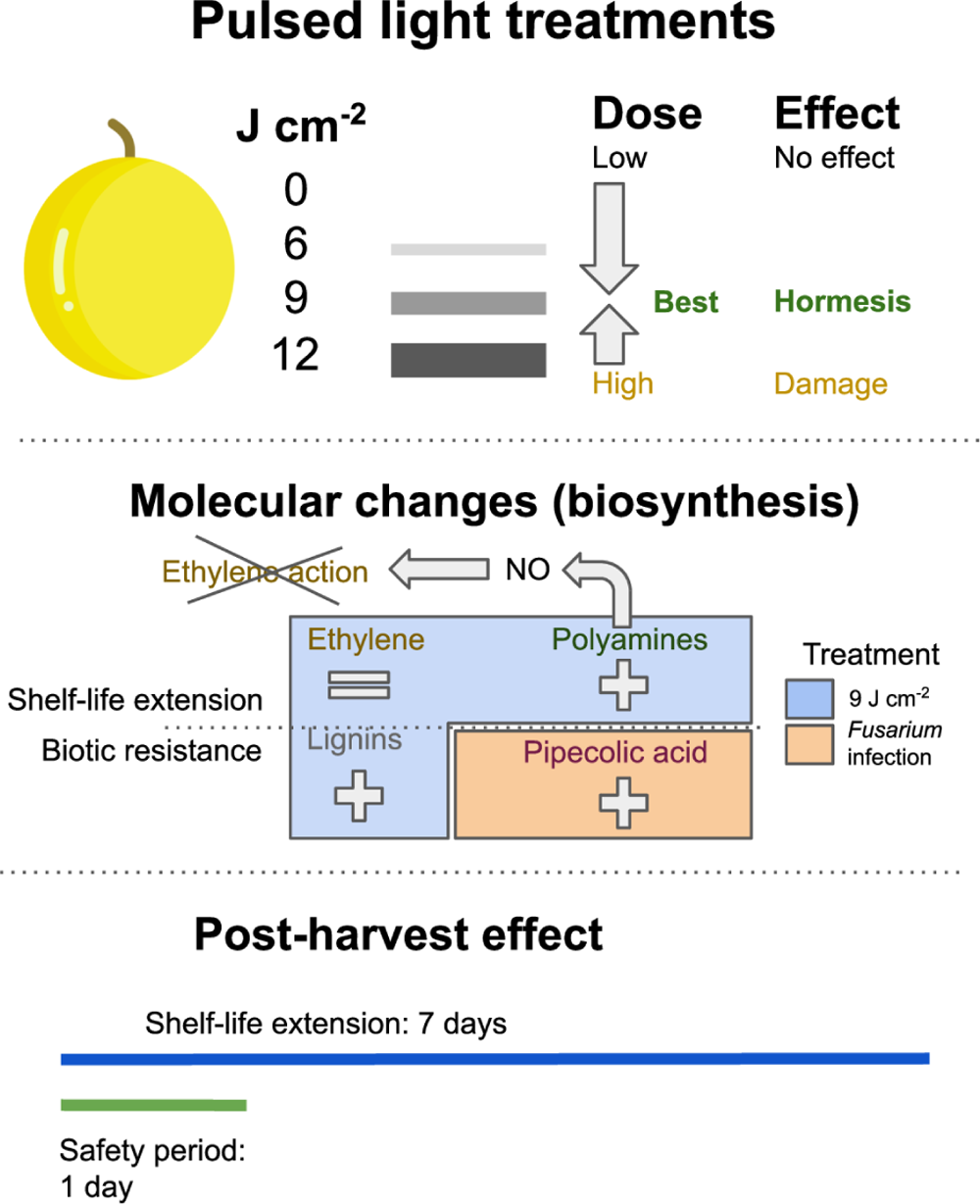
Summary picture of the pulsed light treatment which triggers the hormesis effect on melon fruits.

Another notable advantage of PL treatment was the extension of the shelf life of melon fruits. This positive effect was particularly evident with the same optimal dose that effectively controlled Fusarium infection, resulting in a one-week increase in the fruit shelf life. PL treatments exceeding 9 J cm^−2^ caused significant damage to the fruits, leading to membrane destabilization and loss of firmness (data not shown), which in turn contributed to a shorter shelf life. The observed membrane destabilization, especially in the mitochondrial membrane, disrupts the electrochemical potential and directly impacts energy storage within the cell (Zorova et al., 2018). The high energy content in the UV-C spectrum of PL affects not only DNA but also various other molecules. Excessive UV-C exposure can lead to the disruption of complex molecules like membranes and their integral proteins.

In the case of acerolas (a brazilian berry), blueberries, strawberries and tomatoes, the optimal dose of PL, to control microorganism infection or to extend the shelf life, were identified as 0.6, 1.27, 1.27 and 2.2 J cm^−2^, respectively (Macedo et al., 2023; Salehi, 2022). These berries and tomato, possess a thinner epicarp along with a softer physical barrier, which their optimal PL doses are lower than of melons. In other words, fruits with a less robust epicarp require less energy to induce the hormesis responses. On the contrary, melons, which have a thicker and more substantial epicarp, seem to necessitate a higher energy level to trigger these beneficial responses. The strength of the epicarp thus appears to be a significant factor influencing the energy requirements for inducing hormesis in fruits when applying PL treatment.

One more notable phenotypic observation we made at the 9 J.cm^−2^ dose of PL treatment was an increased level of Fusarium infection occurring at 48h after PL treatment. This phenomenon could be attributed to a metabolic readaptation occurring during this specific time frame of infection. After this critical period, at 96h, it is possible that the fruit returns to a homeostasis stage, which could then trigger a robust immune response against the Fusarium infection. This observation underscores the dynamic and complex nature of the interactions between PL treatment, the fruit metabolism, and the Fusarium infection process. However, physical manipulation of melon fruits after 24 h is not so common in the packing house, guaranteeing in this way a safety period.

### Conclusions

This study provides insights into the transcriptional molecular mechanisms involved in inducing systemic acquired resistance to Fusarium and extending the shelf life of melon fruits through pulsed light treatment. The biological benefits triggered by a 9 J cm^−2^ dose of PL include the upregulation of lignan biosynthesis, enhancing fruit firmness as a physical barrier against fungal infections. Additionally, the upregulation of genes involved in pipecolic acid biosynthesis, when the fruit is infected, plays a crucial role in inducing systemic acquired resistance in plants. These results suggest a dual mechanism in the defense against Fusarium infection in melon fruits promoted by PL and the infection. The extension of fruit shelf life appears to be facilitated by a flux deviation from ethylene biosynthesis to polyamine biosynthesis. In the final step of the polyamine pathway, genes showed upregulation, and one of the subproducts, nitric oxide, is reported to inhibit ethylene biosynthesis and action. Therefore, this study highlights the multifaceted beneficial effects of PL in enhancing the resistance and shelf life of melon fruits.

## MATERIAL & METHODS

### Plant material and *in vivo* inoculation

The study utilized Melon Goldex^Ⓡ^ fruits from Norfruit’s orchards (Lat.: 4°54’15.47”S, Lon.: 37°22’1.86”O) in Mossoró district, Rio Grande do Norte. After harvesting, the fruits were transported to Embrapa Agroindústria Tropical, Fortaleza, CE, on the same day. Upon arrival, the melon fruits underwent washing with sodium hypochlorite (200 μL L^−1^). Subsequently, the fruits were inoculated near the peduncle and five adjacent regions with 100 uL of a suspension containing the *Fusarium pallidoroseum* (106 conidia mL^−1^ - Lima et al., 2021). Following inoculation, the melon fruits were exposed to pulsed light treatments.

### Pulsed light treatment of melon fruits

After 12 hours of the inoculation, when pathogen effects were evident (Cheigh et al., 2012), all fruits underwent PL treatment using the SteriBeam equipment with XeMaticA-2LXL lamps (Germany). The PL doses applied were 0, 6, 9 and 12 J cm^−2^ in a pulsed light chamber. The chamber, equipped with two high power xenon lamps, allowed 360° exposure of the fruits placed on a transparent Teflon^®^ support. Each lamp produced short pulses of 0.3 μs with an energy of 0.3 J cm^−2^ pulse^−1^. The doses were calculated based on the number of pulses: 0 J cm^−2^ (control), 6 J cm^−2^ (20 pulses), 9 J cm^−2^ (30 pulses), and 12 J cm^−2^ (40 pulses). Subsequently, the fruits were incubated in a light-protected box at 28 °C ± 1 °C and 92% relative humidity to allow the mycelial growth. The experimental design consisted of six treatments with three biological replicates each: N00 control (no inoculation, no PL treatment); I20 inoculation with Fusarium and treated with 0 J cm^−2^ PL; I21 inoculation and 6 J cm^−2^ PL; I22 inoculation and 9 J cm^−2^ PL; I23 inoculation and 12 J cm^−2^ PL; and, N22 without infection and treated with 9 J cm^−2^ PL. All PL applications were performed 96 hours post-inoculation. The treatment nomenclature indicates whether there was no inoculation (N) or inoculation (I - first letter), the PL treatment time (0 or 2, indicating 0 hours or 96 hours post-inoculation - first number), and the PL dose applied (last number): 0, 1, 2, 3 (0, 6, 9, 12 J. cm^−2^, respectively).

### RNA isolation and sequencing

After 96 hours of the PL treatment, total RNA extraction was performed. Initially, melon fruits were washed and then dried with toilet paper, and epidermal tissues of the fruits (0.8 cm of diameter and 2.5 cm of thickness) adjacent to the inoculation points were collected for subsequent RNA isolation. The RNA extraction was carried out using CTAB-pBIOZOL protocol (Edmunds, 2017). The quality and quantity of RNA were assessed using both NanoDrop and Agilent 2100 bioanalyzer systems. For RNA-seq library preparation, the TruSeq RNA Library Prep Kit v2 was employed following the manufacturer’s instructions. Subsequently, the libraries were sequenced using the Illumina HiSeq 3000 platform.

### Pre-processing and mapping of reads against the reference genome

The quality of the reads was assessed by the software FastQC Ver. 0.11.6 (Andrews and Others, 2010), before and after the trimming of reads step. Trimmomatic Ver. 0.39 (Bolger et al., 2014) was used to trim bases with low quality and adapters in the reads, with the following parameters: PE, which indicate that the reads are of the type paired-end sequencing; ILLUMINACLIP:TruSeq3-PE.fa:2:30:10 to remove possible adapters; SLIDINGWINDOW:4:15 to remove bases with quality (Q) < 15 in a 4-base window; and, MINLEN:36 to maintain reads with at least 36 bases, discarding any smaller bases after cleaning.

After the trimming step, we performed the mapping of reads against the reference genome of *Cucumis melo* build 3.6.1 and annotation Ver. 4.0 (Ruggieri et al., 2018), obtained from Melonomics database (https://www.melonomics.net/melonomics.html#/). This step was performed by the software Star Ver. 2.5.3a (Dobin et al., 2013) with default parameters. In order to estimate rRNA contamination in the mRNA-seq libraries, we performed an analysis with the software SortMeRNA (Kopylova et al., 2012). To count the reads mapped against the genome we used the software HTSeq Ver. 0.9.1 (Anders et al., 2015). The HTSeq parameters were: counted by gene and no specific strand.

### Identification of differentially expressed genes

The identification of differentially expressed genes (DEGs) was performed with the package edgeR Ver. 3.9 (Robinson et al., 2010) under R environmental Ver. 3.6.1 (Ripley, 2001), based on three libraries (Leading analysis results - Supplementary Material 1). For the identification of DEGs the edgeR cutoff for considering a gene expressed was set to 1 count per million (CPM). After, the data were normalized by the trimmed mean of M values (TMM) method. To consider a DEG, was set as a maximum value of FDR (False Discovery Rate) 0.05.

The comparison of DEG profiles was performed with set theory and graphically represented in Venn diagrams. Moreover, the heatmap and bar graphics were created with ggplot2 Ver. 3.2.1 (Wickham et al., 2008) package in R.

### Biological downstream analysis of the DEGs

In order to characterize function, metabolic pathway, transcription factor (TF) and biological process in each data set of DEGs, the functional annotation from different sources were retrieved. The function annotation of all genes was retrieved from the annotation file obtained in the Melonomics database; genes without annotation were assigned with NA. For the metabolic pathways, the EC (Enzyme Commission) annotated in the GFF3 file were mapped against the KEGG database to reconstruct the metabolic pathways of melon. A TSV (Tab Separated Value) file with all TFs by family was obtained from PlantTFDB Ver. (Jin et al., 2017). The annotation of Gene Ontology (GO) terms by gene was retrieved from melon annotation Ver. 4.0. Due to the massive amount of information, it was performed overrepresentation analysis by type of annotation. To perform this step was applied a Fisher exact test with p-value adjusted by FDR in the annotation class of KEGG pathways and GO terms. An annotation factor was considered as overrepresented under FDR < 0.05, with exception to TF families in which we used p-value < 0.05. Moreover, the representative proteins of each gene locus from the melon genome were submitted to Mercator web server Ver. 3.6 (Lohse et al. 2014) to generate the bins of MapMan Ver. 3.6 (Thimm et al., 2004). The MapMan was used to map melon DEGs in each annotation bin. MapMan uses a kind of annotation that is fast and complementary to KEGG pathways and gene ontology.

### Orthology analysis

The orthology analysis which aims at the identification of orthologous genes between species was carried out using OMA (Orthologous MAtrix) software Ver. 2.5.0 (Train et al., 2017), and the comparisons performed were *C. melo* against *Arabidopsis thaliana* Ver. Araport 11 - 20220914 (Cheng et al., 2017), and *C. melo* against *Solanum lycopersicum* ITAG Ver. 4.0 (Hosmani et al., 2019). To this, representative proteins from each annotated gene were used. Basically, OMA performs local alignment of all proteins between and within the species, in a bidirectional manner to identify high similar groups of proteins.

### Phenotypic evaluation of pulsed light post-treatment effects

Two more experiments, despite the transcriptional evaluation, were done in order to evaluate the pulsed light post-treatment phenotypic effects. The second experiment aimed to assess the residual effect of PL on melon fruits in controlling Fusarium infections. Melon fruits were treated with PL doses of 0, 9 and 12 J cm^−2^. Subsequently, these fruits were inoculated with Fusarium at different time points: 0, 24, 48, and 96 hours after the PL treatment. Each replicate consisted of four melon fruits, for each fruit, four lesions were monitored. The evaluation involved tracking the diameter of the Fusarium infection lesions (measured in cm) from 6 to 18 days after infection, at three-day intervals. Statistical analyses were performed using pairwise comparisons to compare means within hours of infection, and linear regression was also conducted within the same data group. Additionally, pairwise comparisons of infection time were carried out. The statistical analysis was performed in R, utilizing the ggpubr Ver. 0.6.0 package (Kassambara, 2023).

A third experiment was conducted in order to evaluate the extension of fruit shelf life. Survival analysis was performed to control (0 J cm^−2^) and 9 J cm^−2^ dose treatment, each one with 20 replicates. R statistical environment was used to perform the statistical analysis with the packages survminer Ver. 0.4.9 (Kassambara et al., 2021) and survival Ver. 3.5-7 (Therneau, 2024).

## Competing interests

The authors declare that they have no competing interests.

## Funding

This work was supported by the project Tratamento alternativo com luz ultravioleta pulsada no controle de podridões pós-colheita do melão destinado à exportação, code 03.14.16.006.00.00 funded by Embrapa, and Biocomputacional: 30a/2013 - Rede Avançada em Biologia Computacional (RABICÓ), code 88882.160137/2013-01 funded by CAPES.

## CRediT authorship contribution statement

**Luis WP Arge:** Methodology, formal analysis, investigation, writing - original draft. **Guilherme L Morais:** Methodology, writing - review & editing. **Joseane B Carvalho:** Methodology, writing - review & editing. **Guilherme J Zocolo:** Conceptualization, methodology, investigation, writing - review & editing. **Andréia H Oster:** Conceptualization, methodology, investigation, writing - review & editing. **Ana TR Vasconcelos:** Supervision, writing - review & editing. **Leandro E Diniz:** Conceptualization, methodology, investigation, writing - review & editing. **Ebenézer O Silva:** Conceptualization, methodology, investigation, writing - review & editing. **Patricia N Bordallo:** Conceptualization, methodology, investigation, writing - review & editing.

## Declaration of generative AI and AI assisted technologies in the writing process

During the preparation of this work the authors used ChatGPT in order to check the grammar and to improve the readability. After using this tool, the authors reviewed and edited the content as needed and took full responsibility for the content of the publication.

## Data availability

The RNA-seq data underlying this article are available in Gene Expression Omnibus (GEO) Database at https://www.ncbi.nlm.nih.gov/geo/query/acc.cgi?acc=GSE256527, and can be accessed by “GSE256527” identification.

## Notes

### Competing Interest Statement

The authors have declared no competing interest.

